# Multiscale mapping of *in vivo* 3D epidermal melanin distribution of human skin using a fast large-area multiphoton exoscope (FLAME)

**DOI:** 10.1101/2022.02.25.482009

**Authors:** Juvinch R. Vicente, Amanda Durkin, Kristina Shrestha, Mihaela Balu

## Abstract

Melanin plays a significant role in the regulation of epidermal homeostasis and photoprotection of human skin. The assessment of its epidermal distribution and overall content is of great interest due to its involvement in a wide range of physiological and pathological skin processes. Among several spectroscopic and optical imaging methods that have been reported for non-invasive quantification of melanin in human skin, the approach based on the detection of two-photon excited fluorescence lifetime distinguishes itself by enabling selective detection of melanin with sub-cellular resolution, thus facilitating its quantification while also resolving its depth-profile. A key limitation of prior studies on the melanin assessment based on this approach is their inability to account for the skin heterogeneity due to the reduced field of view of the images, which results in high dispersion of the measurement values. Pigmentation in both normal and pathological human skin is highly heterogeneous and its macroscopic quantification is critical for reliable measurements of the epidermal melanin distribution and for capturing melanin-related sensitive dynamic changes as a response to treatment. In this work, we employ a fast large-area multiphoton exoscope (FLAME), recently developed by our group for clinical skin imaging, that has the ability to evaluate the 3D distribution of epidermal melanin content *in vivo* macroscopically (millimeter scale) with microscopic resolution (sub-micron) and rapid acquisition rates (minutes). We demonstrate significant enhancement in the reliability of the melanin density and distribution measurements across Fitzpatrick skin types I to V by capturing the *intra*-subject pigmentation heterogeneity enabled by the large volumetric sampling. We also demonstrate the potential of this approach to provide consistent measurement results when imaging the same skin area at different times. These advances are critical for clinical and research applications related to monitoring pigment modulation as a response to therapies against pigmentary skin disorders, skin aging, as well as skin cancers.

## INTRODUCTION

Melanin is a group of molecules with multifunctional characteristics and is the most abundant chromophore in the human skin. The prevalent forms of melanin in the skin are eumelanin (brown-black) and pheomelanin (yellow-reddish). They are produced by specialized cells, called melanocytes, mainly found in the basal layer of the skin epidermis.^1–3^ Stored in melanosomes, they are transferred to keratinocytes to protect the skin from photo-damage against UV radiation.^4^ Melanin plays a significant role in a variety of physiological and pathological conditions. Reliable measurements of the cutaneous melanin distribution and the melanin-related sensitive dynamic changes are essential for a better understanding and more efficient treatment of pigmentary skin disorders and for differentiating melanoma from benign pigmented lesions.

The current gold standard methods for melanin quantification in skin involve *ex vivo* chemical analyses.^5^ These approaches are invasive and thus, impractical to be performed repeatedly for applications such as those related to pigmentary skin disorders that require monitoring of the treatment response.^6^ They are also not feasible for the assessment of the melanocytes density, a key metric in quantitative approaches proposed for the non-invasive diagnosis of melanoma.^7^ Optical methods based on reflectance spectroscopy and colorimetry have been developed to map the distribution of melanin in human skin *in vivo*.^8–11^ However, the epidermal depth-information and spatial resolution provided by these approaches are limited. Multispectral photoacoustic imaging enables non-invasive 3D mapping of epidermal melanin based on the detection of ultrasonic waves generated by rapid thermoelastic expansion of melanin induced by pulsed laser irradiation.^12–14^ Remarkably, this approach generates images of melanin and blood vessels from 1 to 2 mm below the skin surface, but the 5-30 μm spatial resolution of the images is not sufficient for resolving the epidermal melanin depth profiles.^13,14^ In recent work, Yakimov et al. reported on a promising approach for determining epidermal melanin depth-distribution based on fluorescence and Raman spectroscopy data acquired from two subjects with different skin types.^15^ An approach that has been demonstrated to be effective for the quantification of epidermal melanin, while also resolving its depth profile, is based on the detection of melanin two-photon excited fluorescence (TPEF).^6,16–18^ TPEF laser-scanning microscopy technologies can generate *in vivo* 3D sub-micron resolution images of epidermal melanin distribution in human skin.^6,16–19^ The penetration depth of this imaging technique is limited to 150-200 μm depending on the skin type and other imaging parameters.^20,21^ Nonetheless, this depth is sufficient for capturing the entire thickness of the epidermis on most areas of the body.^22^ Enhanced specificity of the melanin detection in the skin is further achieved using two-photon fluorescence lifetime imaging (FLIM) by exploiting the melanin fast fluorescence decay (<0.2 ns) compared to the fluorescence decay of other fluorophores in the skin.^18,19,23^ A. M. Pena *et al*. demonstrated the feasibility of this approach by studying the modulation of the 3D epidermal melanin following long-term topical treatments.^17,18^ FLIM, among other methods such as pump-probe^24^ and Coherent anti-Stokes Raman Scattering imaging,^25^ has also been demonstrated as a potentially effective tool for selective detection of eu- and pheo-melanin in excised pigmented lesions from human skin and *in vivo* in murine model^24,25^ and human skin.^23^ To enhance the clinical imaging feasibility of this approach, our group and others have proposed and demonstrated significant enhancement in image acquisition time based on temporal binning^26^ along with analysis of the fluorescence temporal decay slope.^17,18^

While the potential of the two-photon FLIM approach for quantification of melanin and its depth-distribution in human skin has been demonstrated in the clinical setting, a key limitation remains. This limitation is related to the reduced field of view (0.25×0.25 mm^2^) of the commercial clinical multiphoton tomograph^27,28^ that has been used in prior studies for evaluating the potential of the TPEF intensity^6,16^ and lifetime^17–19^ detection in the assessment of melanin in human skin. Given the heterogeneous nature of both normal and pathological skin, generating depth-resolved images over wide areas along the skin surface is critical for reliable measurements of the epidermal melanin distribution and of the melanin-related sensitive dynamic changes as a response to treatment.

Recently, our group has developed a fast large-area multiphoton exoscope (FLAME),^26^ a multiphoton imaging system optimized for clinical skin imaging that has the ability to rapidly generate 3D images (within minutes) over macroscopic areas of skin (up to 1 cm^2^) with sub-cellular resolution (0.5-1 μm). Combined with fluorescence temporal gating and binning for melanin-specific detection, this system is capable of quantifying melanin almost in real-time. In this work, we employed FLAME to evaluate *in vivo*, in human skin, the significance of the increased imaging area on the epidermal melanin measurement’s reliability.

## RESULTS

### Macroscopic mapping of melanin volume fraction for different Fitzpatrick skin types

We employed FLAME to measure the melanin density in human skin based on volumetric multiphoton microscopy (MPM) images acquired from the volar and dorsal forearms of subjects with Fitzpatrick skin types ranging from I to V (Figure 1). Figure 1a shows the macroscopic maps (millimeter scale) for the melanin volume fraction (MVF) as *z*-projections corresponding to sun-exposed (dorsal forearm) and non-sun-exposed (volar forearm) areas for subjects with skin types I-V. These images clearly illustrate the heterogeneity in the melanin distribution, particularly for skin type I. To evaluate the *z*-distribution of the epidermal melanin content, we measured the average melanin density for each epidermal layer. Here, the position across the epidermis was normalized against the epidermal thickness starting from the basal layer (0) to the top of the epidermis (1) (Figure 1b). We generally measured a higher melanin density in the epidermal layers close to the *stratum basale* compared to the upper epidermal layers. For the volar forearm of subjects with skin types I-IV, the epidermal melanin *z-*distribution showed mostly a lack of pigmentation in the upper third of the epidermis. In the dorsal forearm of the same subjects, the melanin was present across the entire epidermis with higher values in the layers close to the *stratum basale*. The skin type V subjects showed elevated melanin content across the entire epidermis for both their volar and dorsal forearms compared to subjects with lighter skin types. The global MVF values for the subjects’ dorsal and volar forearms are summarized in Figure 1c. As expected, the global MVF values increase with skin type and exhibit an overall change of ∼7× from skin type I to V. Although not linear, this increasing trend in MVF correlates strongly with the lightness (*L*\*) parameter from the colorimetry measurements (Supplementary Figure S1). One-way analysis of variance (ANOVA) shows a significant difference in the average MVF values corresponding to both the volar and dorsal forearms among all skin types, except for skin types II and III (Supplementary Tables S1). Lastly, the average MVF values between the volar and dorsal forearms also show a significant difference for skin types II-V (Supplementary Table S3). This is consistent with the lower *L\** parameters measured from the dorsal forearm compared to the volar forearm (Supplementary Figure S1).

**Figure 1.**
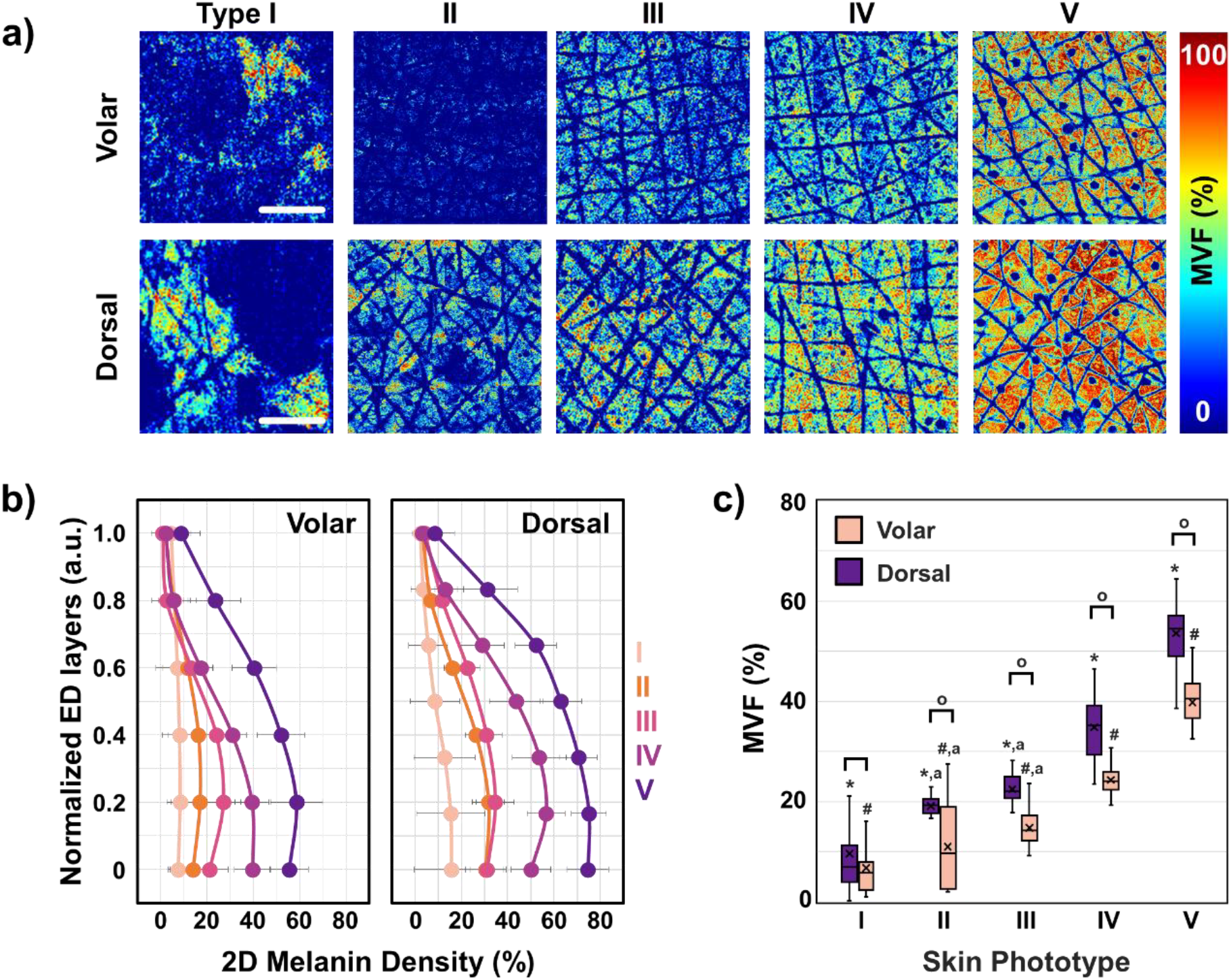
3D melanin density for Fitzpatrick skin types I-V. (a) The *z*-projection of MVF for representative subjects with skin types I-V, from the volar and dorsal forearm. Scale bar = 1 mm. (b) 2D-melanin density as a function of epidermal depth from the basal layer (0) to the top of the epidermis (1) for all the skin types. The data and error bars represent the average and standard deviation of the melanin fraction in each layer, respectively. The position across the y-axis is normalized against the epidermal thickness. (c) The global MVF values for all the skin types from both dorsal and volar forearms. (*, #) indicate a significant difference (*P*<0.01) among the average MVF values in the dorsal and volar forearm, respectively, except for skin types sharing letter ‘a’. (^O^) indicate a significant difference (*P*<0.01) between dorsal and volar forearm within a skin type. (*N* = 32)

### Macroscale mapping of melanin volume fraction captures intra-subject heterogeneity

While the MVF values and the 2D melanin density profiles across the epidermis were determined based on the total melanin content imaged at different depths over a skin area of 3.2×3.2 mm^2^, a closer examination of the melanin distribution maps reveals the variability of these values along the epidermis because of the skin heterogeneity. The images and data in Figure 2 illustrate the degree of variability of the melanin density along and across the epidermis within representative subjects with light and dark skin types I and V, respectively. Thus, the skin type I subject (Figure 2a) presents skin areas characterized by MVF values as low as 0.3% (ROI-1) in close proximity to areas characterized by MVF values as high as 52% (ROI-3). The corresponding melanin density *z*-depth profiles (Figure 2b) show a very distinct distribution for each ROI, where the ROI-3 has high melanin content in all epidermal layers while the ROI-1 is almost void of it across the entire epidermis. The skin type V subject presents a more uniform skin pigmentation when visualized macroscopically (Figure 2c), yet at the microscopic scale, the MVF values show some degree of heterogeneous pigmentation with areas of 78% MVF (ROI-3) in close proximity of areas with 52% MVF (ROI-1). The areas characterized by high MVF values (ROI-3) showed a higher melanin content in the upper epidermal layers compared to skin areas with lower MVF values (ROI-1 and ROI-2) as illustrated by the melanin density *z*-distribution profiles (Figure 2d). Note that we analyzed the MVF values and corresponding 2D melanin density profiles over regions of interest (ROIs) of 0.25×0.25 mm^2^. The rationale for emphasizing the skin heterogeneity when examining this area size is related to the fact that this is the skin area typically scanned with the current commercial clinical multiphoton tomograph employed in prior studies for *in vivo* melanin quantification in human skin.^16–18^

**Figure 2.**
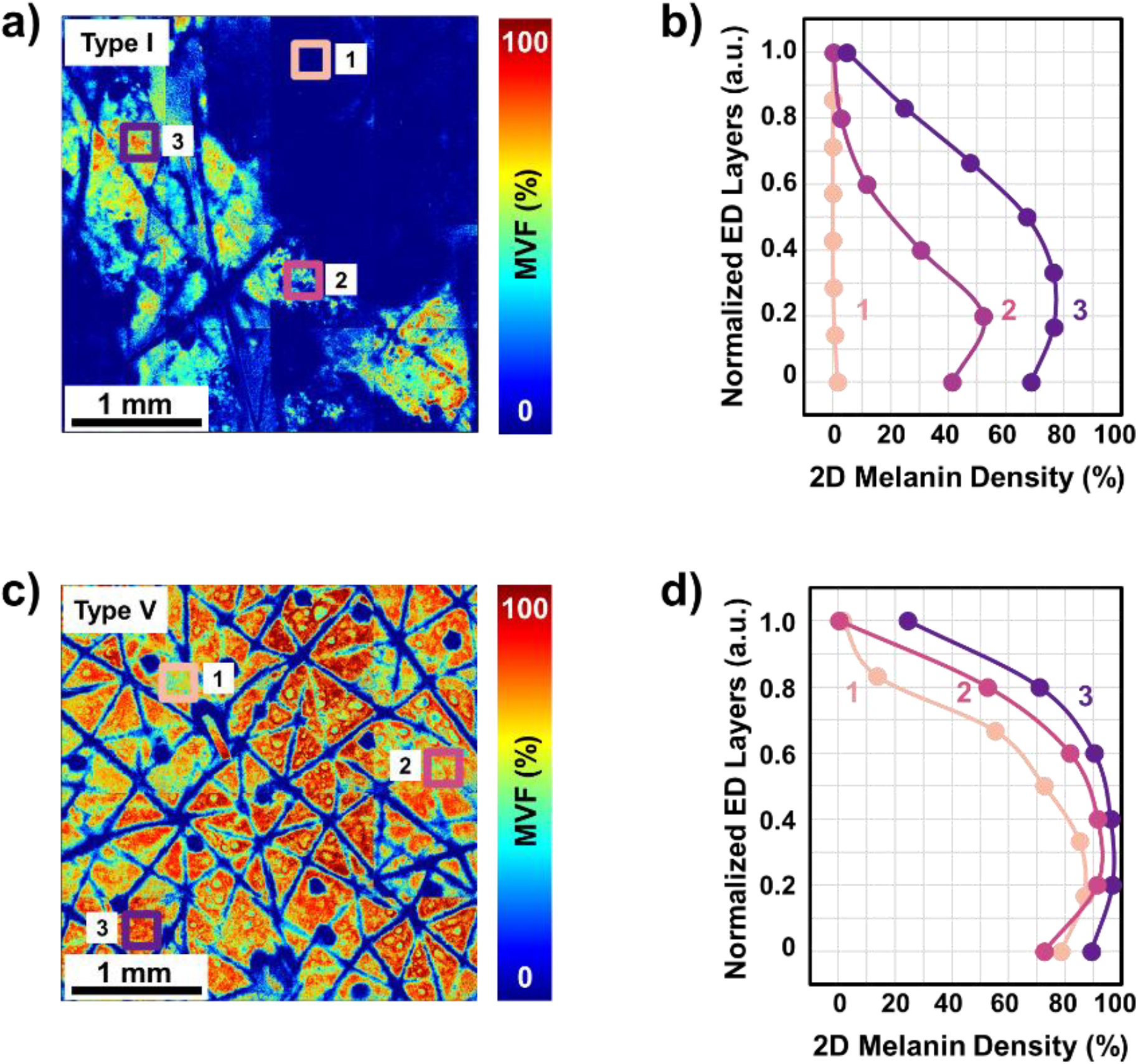
Comparison of 3D melanin density for skin types with different levels of heterogeneity. (a, c) *z*-projection of the MVF values from the dorsal forearm of representative subjects with skin types I and V, respectively. (b, d) 2D melanin density as a function of epidermal depth from the basal layer (0) to the top of the epidermis (1) for skin types I and V, respectively. The position across the y-axis is normalized against the epidermal thickness. The plots in (b) and (d) correspond to the ROIs of 0.25×0.25 mm^2^ marked in (a) and (c) with the same color and number.

### The precision of global MVF measurements increases with an increase in the imaging area

The results described above show the heterogeneity in the 3D distribution of melanin in human skin for all skin types, which further suggests that a large sampling volume is critical for the reliability of the MVF measurements as they would capture this heterogeneity. To test this hypothesis, we evaluated the MVF values corresponding to different imaging areas. We performed these measurements by dividing the 3D stacks acquired over the entire scanning area of 3.2×3.2 mm^2^ into 3D stacks of sub-images with sizes ranging from 0.25×0.25 to 1.6×1.6 mm^2^ (see *Methods* for more details). The MVF values were calculated for all the sub-images in the 3D stacks. The corresponding average and standard deviation (S.D.) values are summarized in Figure 3. Our measurements showed the averages of the MVF values are independent of the size of the imaging area for all skin types provided that for small scanning areas a large number of stacks is acquired to encompass a macroscopic, mm-scale area. However, while this approach can estimate the average MVF value, it does so with a large error margin. Based on the data presented in Figure 3c-d, the variation in the MVF values, based on their S.D. values, decreases with the increase in the imaging area. Our measurements also showed a rapid decrease in the standard deviation of the MVF values for the small range of imaging areas (0.25×0.25 to 0.65×0.65 mm^2^), followed by a more gradual change for the relatively large imaging areas (0.80×0.80 to 1.6×1.6 mm^2^). An increase in the FOV size from 0.25×0.25 mm^2^ to 1.6×1.6 mm^2^ results in an average of ∼2-fold and ∼3-fold increase in the MVF precision for the volar and dorsal forearms, respectively, across the skin types.

**Figure 3.**
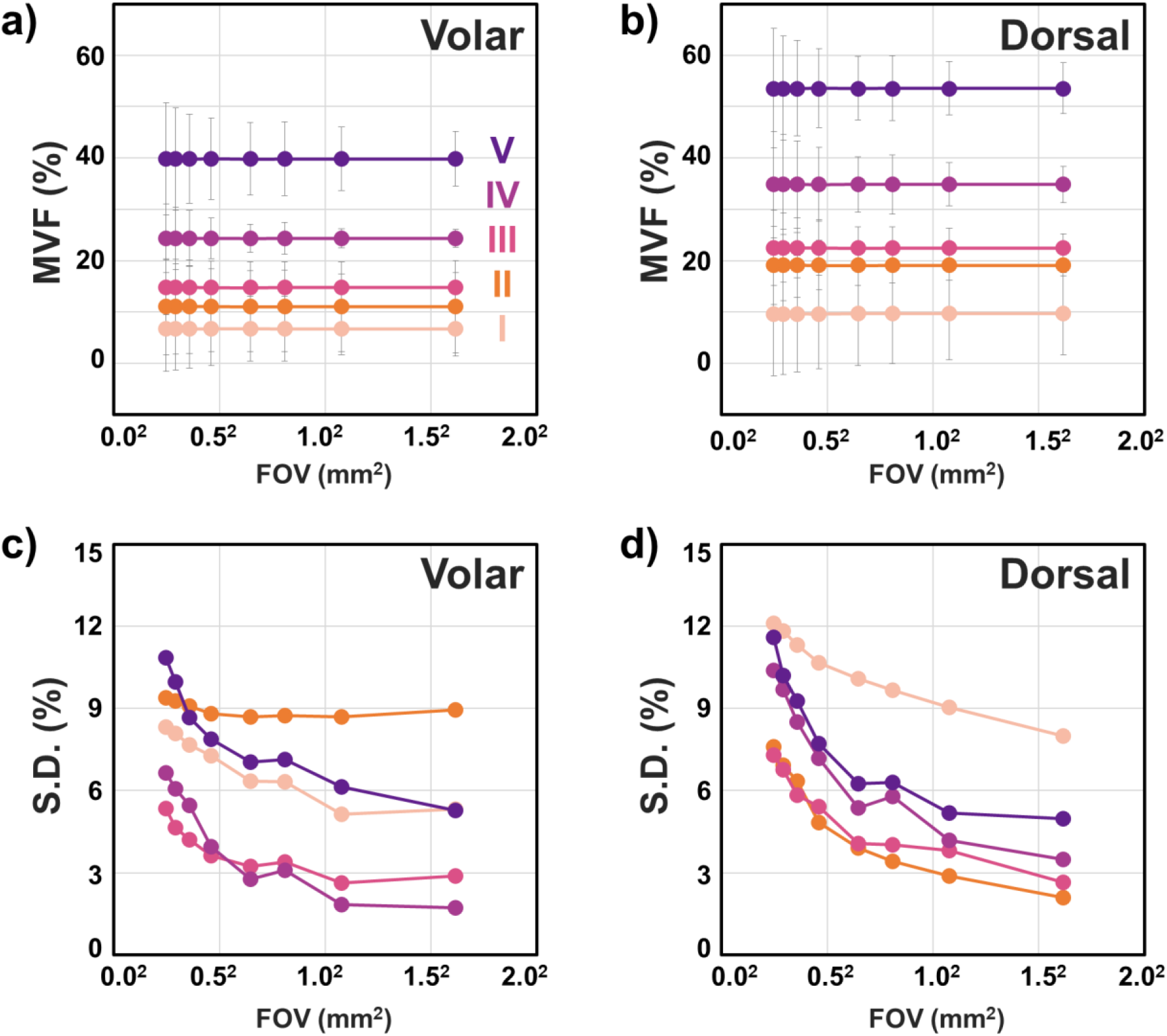
The effect of field of view on the accuracy and precision of the MVF measurements. (a, b) The MVF for skin types I to V for the volar and dorsal forearm, respectively. The data points and error bars correspond to the average MVF and their standard deviation (S.D.) (c, d) The S.D. of the MVF values as a function of imaging area for skin types I to V for the volar and dorsal forearm, respectively. (0.25×0.25 mm^2^, *N* = 338, 1.6×1.6 mm, *N* = 8. Complete list in *Methods*).

### Large-area sampling results in more precise MVF measurements compared to selective small-area sampling

As illustrated in the images above, the MVF maps measured over macroscopic, mm-scale skin areas include skin folds, hair follicles, and pores. These features are too large for measurements performed over smaller, sub-mm scale scanning areas. In this case, the images are acquired such that they are void of these large features as reported in the prior studies employing the current clinical multiphoton tomograph that generates images with a FOV limited to 0.25×0.25 mm^2^.^6,16,17^ To determine the effect of this subjective approach on the MVF measurements we compared the MVF results obtained by two methods: Method 1) manual sampling of 0.25×0.25 mm^2^ ROIs (*N* = 9) from the 3D stacks acquired over the entire 3.2×3.2 mm^2^ scanning area such that the selected ROIs were void of the large features mentioned above and Method 2) objective sampling by dividing the entire scanning area of 3.2×3.2 mm^2^ into 1.1×1.1 mm^2^ ROIs (N=9). The results obtained from three representative subjects with different skin types are summarized in Figure 4. For all subjects, the large-area, objective sampling approach (Method 2) generally resulted in significantly less variation of the MVF compared to the values obtained by selectively scanning small-areas (Method 1). Thus, the representative examples presented in Figure 4, show a 2-fold (S.D. 6% → 3%) and 3-fold (S.D.: 9% to 3%) increase in the precision of the MVF measured in the volar and dorsal forearm, respectively of the skin type I subject by using the objective sampling (Method 2) (Figure 4c). Similarly, the same approach resulted in about 2-fold enhancement of the MVF precision (S.D.: 7% to 3%) for the values measured in both volar and dorsal forearm of a skin type V (Figure 4e). An additional finding is related to generally higher MVF average values obtained by using the selective scanning-area sampling (Method 1) compared to the large scanning area approach (Method 2). This is associated with the exclusion of the skin folds, hair follicles, and pores in the approach of Method 1. These features are generally devoid of melanin, and their exclusion results in an overestimation of the MVF values.

**Figure 4.**
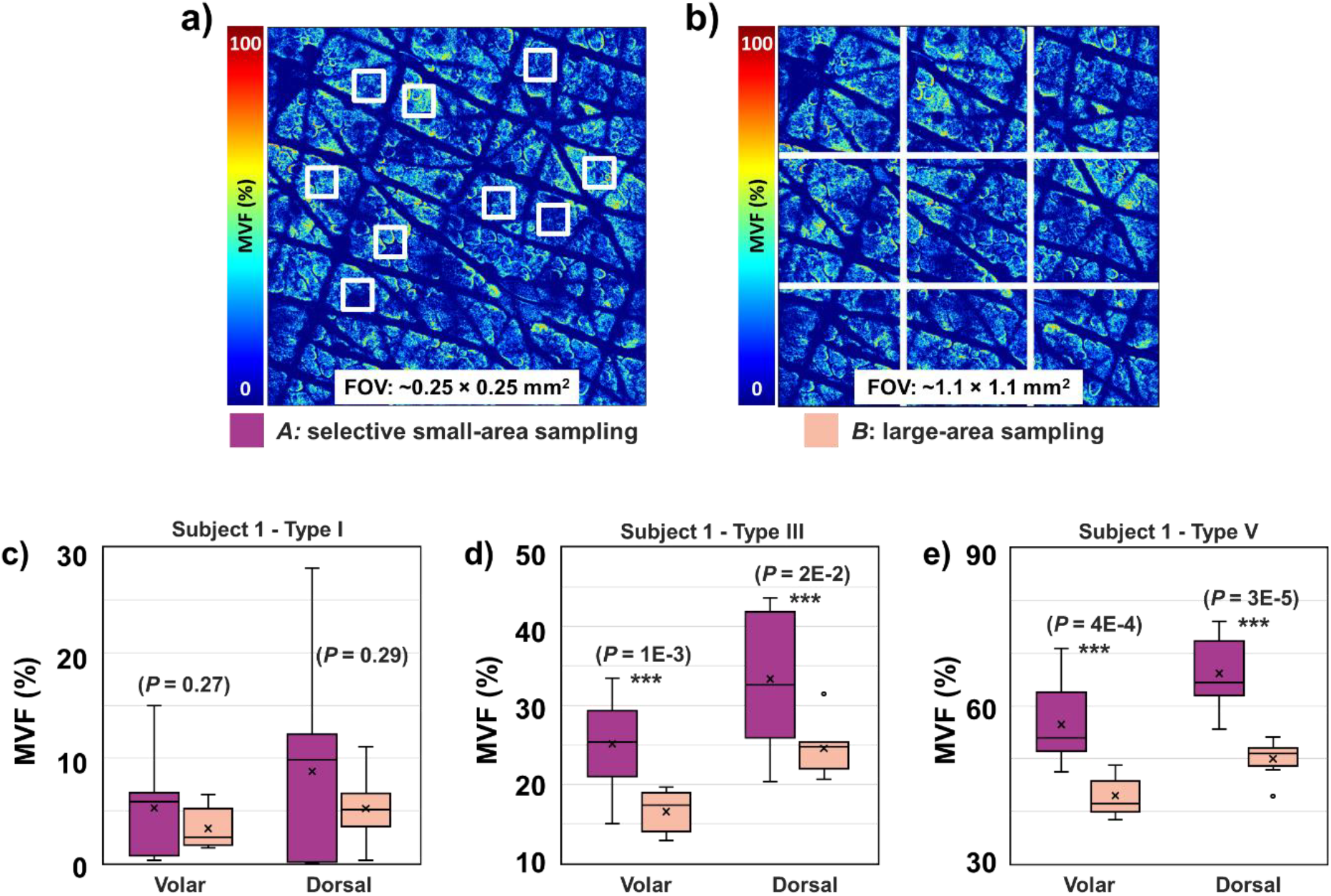
Comparison of the MVF measured by selectively sampling a small-area (0.25×0.25 mm^2^) *versus* sampling a large-area (1.1×1.1 mm^2^). (a, b) Schematic diagrams for Methods *A* and *B*, respectively. (c, d, e) The resulting MVF values using the sampling methods described in (a) and (b) for skin types I, III, and V, respectively (*N* = 9). (***) Indicate significant difference between the MVF values obtained using Method *A* and *B* using a two-sample unpaired t-test with resulting *P*-values indicated in the figure.

### The MVF values are consistent between measurements

Besides accuracy and precision, reproducibility is another parameter of interest for the melanin density measurements in the skin. While the reproducibility assessment requires a dedicated rigorous study on a larger number of subjects, we aimed to provide a brief insight into the potential of our proposed approach to generate consistent results for the melanin density measurements. Thus, we performed MVF measurements on a skin type IV subject’s dorsal forearm at three different times. We used skin landmarks to ensure imaging of approximately the same location. The results are summarized in Figure 5. One-way ANOVA shows no significant difference among the average MVF measurements (*P* = 0.432, Supplementary Table S4).

**Figure 5.**
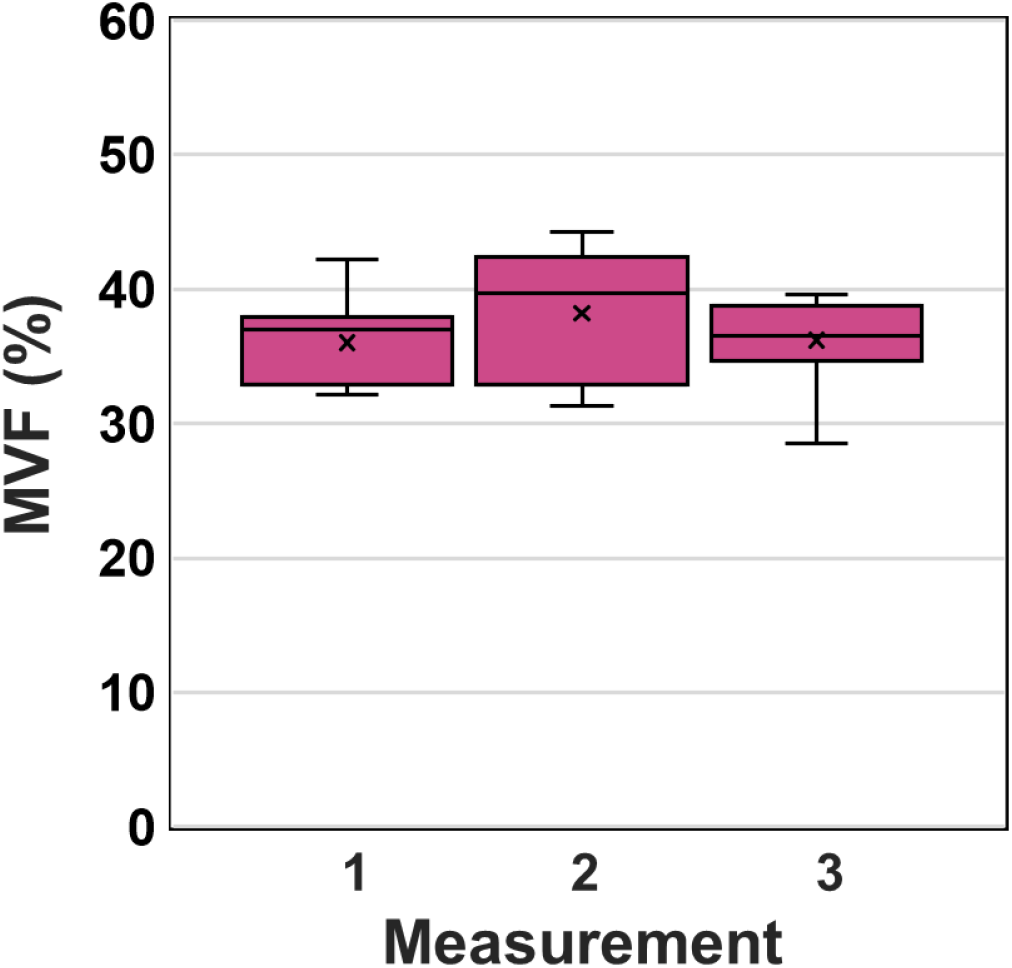
Repeatability of the MVF measurements. MVF values measured at three different times by imaging the dorsal forearm of a skin type IV subject. (*N* = 9, FOV: 1.1×1.1 mm^2^ for each measurement). All measurements were performed based on images acquired from approximately the same area using skin landmarks as guidance.

## DISCUSSION

The distribution of melanin in human skin is innately heterogeneous. In this work, we utilized FLAME, a fast large-area multiphoton exoscope to evaluate the 3D melanin distribution over an imaging area of about two orders of magnitude (180×) larger than the 0.25×0.25 mm^2^ FOV, commonly used for the melanin assessment in human skin using the commercial clinical multiphoton imaging technology.^16–18^ Combined with selective detection of melanin by fluorescence temporal gating and binning, the data acquisition time for each subject was kept under ∼7 minutes, which is at least an order of magnitude faster than the acquisition time that would be required for the current MPM clinical technology to scan over a similar area.

We validated the ability of FLAME to assess the melanin density and distribution in human skin by demonstrating the correlation of the global 3D epidermal melanin density with the skin type and its distribution across the epidermis. The different skin types’ skin areas were characterized by different MVF values with only skin types II and III showing a non-significant difference based on the MVF values for both the sun-exposed (dorsal forearm) and non-sun-exposed (volar forearm). This was likely due to inaccuracy in the determination of the skin type, a hypothesis supported by the colorimetry measurements (Supplementary Figure S1). Correlations of the MVF values with skin type have been reported in prior studies based on TPEF intensity^16^ and lifetime^18^ imaging of melanin, but the statistical significance of the difference in MVF values among different skin types has not been evaluated or reported. One study employing the TPEF lifetime for cutaneous melanin quantification reported on limitation in resolving the melanin concentration in light skin types and in distinguishing the melanin concentration between volar and dorsal forearm due to insufficient precision of the measurements.^19^

The main findings of our work are related to the capability of FLAME to capture the pigmentation heterogeneity in human skin, which has several key implications for studies that involve monitoring melanin modulation as a response to therapies. One implication is that the epidermal melanin density measurements derived from 3D images acquired over macroscopic (mm scale) skin areas are significantly less affected by variability than the measurements obtained from volumetric images acquired over multiple small areas (sub-mm). Technically, the average melanin density values are independent of the scanning area for all skin types and are the same regardless of the sampling approach: rapidly over fewer large areas or slower over more small areas, but the precision of the measurement is higher in the former case. However, in practice, when using the current commercial clinical MPM devices (DermaInspect or MPTflex, Jenlab, Germany) the average melanin density values may be overestimated when compared to the data derived from macroscopic imaging, according to our analysis. The reason is related to the approach used for the 3D melanin density measurements derived from images acquired with the commercial MPM clinical devices where the scanning area (< 250×250 μm^2^) is selected such that it would not include large features such as skin folds, hair follicles or pores. These features are generally devoid of melanin and do not have a significant contribution to the measured melanin content, but they do contribute to the overall volume when scanning macroscopic areas.

The enhancement in the measurement precision of melanin density and distribution is expected to require fewer images to detect certain levels of variations in the MVF values in the skin. An example of a sample size estimate as a function of FOV is shown in Supplementary Table S5 based on the data acquired from the dorsal forearm of the skin type V subjects. As expected, employing the largest FOV (1.6×1.6 mm^2^) would require the smallest sample size for detecting 10% to 25% changes in the MVF values. To detect the same change, the smallest FOV (0.25×0.25 mm^2^) requires ∼4 to 5 times higher sample size. The fewer samples required when using a larger FOV for imaging could translate to a further decrease in the acquisition time. These results were based on the limited data acquired from the dorsal forearm of the skin type V subjects, but we expect a similar trend for all skin types.

The reproducibility of the measurements is a key measurement parameter of interest in clinical studies besides reliability. We performed a brief evaluation of this metric by imaging the forearm skin of a subject at three different times in approximately the same location. The data demonstrated the potential of FLAME to generate images that would result in consistent measurements of the 3D epidermal melanin density. We attribute the consistency of the results to the insensitivity of the large-area sampling to the accuracy of the location selected for imaging. This is particularly of interest in monitoring pigment modulation as a response to therapy or to cosmetic products, where changes can be subtle, and locating the same imaging area for performing the measurements at different time points may be a challenge. A statistically powered study on a larger number of subjects will be required for a rigorous assessment of the melanin density measurements reproducibility.

One important benefit of this approach is the ability to image the melanin with high specificity based on the melanin’s fast fluorescence decay compared to the fluorescence decay of other endogenous fluorophores in the skin. The images are generated rapidly by binning the time-gated TPEF signals rather than by using conventional approaches based on fitting of the full fluorescence decay or phasor analysis, which are not feasible for clinical imaging due to their long integration times. A drawback of the detection based on the time binning of the fluorescence signal is related to the sensitivity of classifying the pixels corresponding to melanin in the TPEF images with respect to the selected threshold. In a recent publication, Pena *et al*. performed a thorough analysis to establish a threshold for melanin detection using a method based on temporal binning and fluorescence temporal decay slope analysis, referred to as pseudo-FLIM. Although our approach is slightly different in several aspects, the mean melanin density values are comparable with the values derived in Pena’s publication for all skin types granted the comparison is a relatively rough estimation due to the different error margins, significantly lower for our measurements.

## CONCLUSION

In this study, we employ FLAME, a fast, large area multiphoton exoscope, a device with unique performance features optimized for clinical skin imaging, to evaluate the significance of the large volumetric sampling enabled by this instrument on the cutaneous melanin measurements reliability. We demonstrated significant enhancement of the melanin density and distribution measurements reliability across Fitzpatrick skin types I to V based on the ability of this imaging device to capture the intra-subject pigment heterogeneity at the microscopic scale. We also demonstrated the potential of this approach to provide consistent measurement results when imaging the same skin area at different times. We expect these advances to have an important impact in clinical and research applications related to monitoring pigment modulation as a response to therapies against pigmentary skin disorders, skin aging, as well as skin cancers.

## METHODS

### Experimental design

We enrolled in the study a total of ten (10) subjects with Fitzpatrick skin types I-V, two (2) for each skin type. We acquired the *in vivo* MPM images from the skin areas on the dorsal and volar left forearm of the subjects. The experiments were conducted with full consent from each subject using an approved protocol by the Internal Review Board for clinical research in human subjects at the UC Irvine.

### Clinical multiphoton imaging

#### Fast large-area multiphoton exoscope (FLAME)

All multiphoton imaging was performed using the clinical Fast Large-Area Multiphoton Exoscope (FLAME), recently developed by our group.^26^ Briefly, this clinical MPM imaging system consists of a turn-key femtosecond laser (Carmel 780, <90 fs, 80 MHz, fixed 780 nm excitation, Calmar, Palo Alto, CA), an articulated arm attached to the imaging head housing near-infrared (NIR) optics: a 4kHz resonant-galvo beam scanning module, custom-designed relay, beam-expander optics and a 25×, 1.05 NA objective lens (XLPL25XWMP, Olympus). The optical design of this system is optimized to provide sub-micron resolution images with a single field of view of up to ∼0.8 × 0.8 mm^2^ at a rapid rate of 7.5 frames per second for ∼1024 × 1024 pixels frame. The imaging area can be increased to ∼10 × 10 mm^2^ by using the tile or strip mosaic approach where adjacent field-of-views are stitched together as described in detail in our report on this device.^26^ The tile mosaic approach was used in this study. The rapidly acquired images are post-processed using a deep-learning-based algorithm, CARE (Content-Aware Image Restoration),^29^ trained specifically for MPM images of human skin as previously described.^26^

The system has two (2) hybrid-photomultiplier tube detectors utilized for simultaneous acquisition of second-harmonic generation (SHG) (330 – 480 nm), and TPEF signals (510 – 610 nm). We further split the TPEF signal into two-coarse components based on the lifetimes of the fluorescence signal using temporal gating. Thus, each image is generated by simultaneously collecting the signals from three (3) separate channels in real-time. Channel-1: the second harmonic generation (SHG, blue channel) signal mainly from collagen, channel-2: short lifetime TPEF (0-1.6 ns, red channel), consists mainly of photon counts from melanin, and lastly, channel-3: long lifetime TPEF (1.6-12.5 ns, green channel). An example of an acquired image is shown in Supplementary Figure S2.

#### In vivo 3D multiphoton imaging

For each subject, we acquired 4×4 mosaic images from the volar and dorsal forearm, encompassing a total area of ∼3.2 × 3.2 mm^2^ (0.84× 0.84 mm^2^, 1024 × 1024 pixels per tile, 10 frames accumulation, 2% overlap per tile). To fully capture the depth of the epidermis, a 3D *z*-stack was acquired for each tile, from a depth slightly above the *stratum corneum* to the dermis (12 optical sections at 10 µm *z*-step). The typical acquisition time for each mosaic image is ∼6.5 minutes (24 s per 3D *z-*stack). The images are post-processed immediately after imaging by using the CARE denoising algorithm. *Laser power considerations:* Our FLAME system uses an objective of NA = 1.05 and an excitation laser power of 60 mW at the skin surface, which is below the DNA and thermal damage threshold limits established for two-photon microscopy of human skin^30,31^; notably, FLAME uses a significantly lower laser fluence (1.7×) and faster imaging time per unit area (40×) compared to the values used in the establishing the damage threshold for two-photon microscopy of human skin.^30,31^

### Colorimetry measurements

The pigmentation in both dorsal and volar regions of the subjects’ forearm was also measured by reflective colorimetry using a CR-400 Colorimeter (Konica Minolta Sensing Inc., Japan) calibrated to black and white calibration plates. The reflected light is collected for a tristimulus color analysis using the *L*\**a*\**b* color system as determined by the *Commission Internationale de l’éclairage* (CIE). The measurements were done immediately after the MPM imaging on around the same imaging spot.

### Data Analyses

#### Melanin content quantification

The melanin content for each image was quantified as previously described.^26^ Briefly, a binary melanin mask is generated by subtracting the photon counts of the green channel from the red channel. All pixels with differences in photon counts > 0 are set to “1”. We then apply an open area filter to remove isolated single pixels which correspond to melanosome areas <1µm^2^. The resulting mask represents the fraction of the total area occupied by melanin in each image. Similarly, these pixels represent the melanin volume fraction (MVF) occupied by melanin in each 3D *z*-stacks. The scheme for this approach is shown in Supplementary Figure S4.

#### Effect of FOV size on MVF variability

To estimate the variation in the MVF measurements as a function of the imaging FOV size, the 3.2×3.2 mm^2^ mosaic 3D stack was divided into sub-images with sizes 0.25×0.25 mm^2^ (*N*=169), 0.29×0.29 mm^2^ (*N*=121), 0.36×0.36 mm^2^ (*N*=81), 0.46×0.46 mm^2^ (*N*=49), 0.65×0.65 mm^2^ (*N*=25), 0.81×0.81 mm^2^ (*N*=16), 1.08×1.08 mm^2^ (*N*=9) and 1.6×1.6 mm^2^ (*N*=4).

#### Selective small-area samples vs large-area samples

Method A: Nine (9) ROIs with 0.25×0.25 mm^2^ size were manually selected from a representative mosaic image to avoid the inclusion of large features such as hairs, skin folds, and pores. Method B: The full mosaic 3D stack was divided into 9 ROIs with 1.1×1.1 mm^2^ size. In both methods, the average MVF and standard deviation were obtained.

#### Measurement repeatability

To assess the repeatability of the MVF quantification, we did multiple measurements on a representative subject (skin type IV, dorsal forearm). Skin landmarks were used to ensure imaging of approximately the same location.

#### Statistical analysis

To compare the average MVF values between dorsal and volar forearms, and between selective small-area sampling and large-area sampling (Figure 4), we performed a two-sample unpaired t-test to determine the significant difference. The significance level threshold is set at *P* < 0.05. For the global MVF for each skin type from dorsal and volar forearms (Figure 1) and the repeatability of the measurement (Figure 5), the average MVF values obtained were compared by one-way analysis of variance (ANOVA) with Tukey-Kramer posthoc analysis. The significance level threshold was set at *P* < 0.05. The sample size analysis in Table 1 is based on a two-sample unpaired t-test (power: 80% and significance level: 5%). Effect size (*d*) was estimated for detection of 10% to 25% change in the mean MVF values using different FOV sizes.

## DATA AVAILABILITY

The datasets generated and/or analyzed in this study are available from the corresponding author upon request.

## ACKNOWLEDGEMENTS

This work was supported by the National Institute of Biomedical Imaging and Bioengineering (R01EB026705) and the UCI Skin Biology Resource-Based Center (P30AR075047). The content is solely the responsibility of the authors and does not necessarily represent the official views of the National Institute of Health.

## AUTHOR INFORMATION

### Contributions

Conceptualization and study design: J.R.V, M.B. Optimization of the instrumentation for *in vivo* imaging: A.D. Data collection: J.R.V, K.S. Development of methodology: JRV, M.B. Data analysis and interpretation of data: J.R.V., M.B. Manuscript writing: J.R.V and M.B. Manuscript review: all authors.

