## Supplementary Data for "Multiscale mapping of *in vivo* 3D epidermal melanin distribution of human skin using a fast large-area multiphoton exoscope (FLAME)"

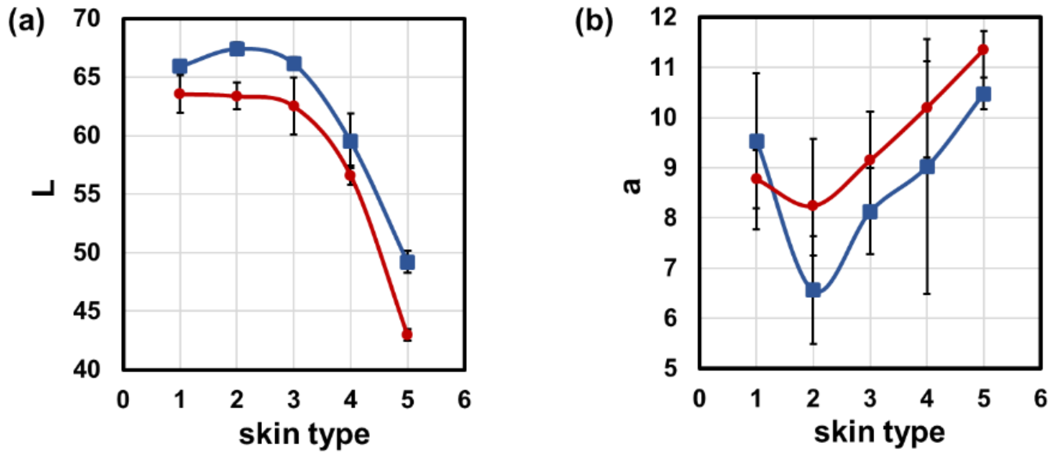


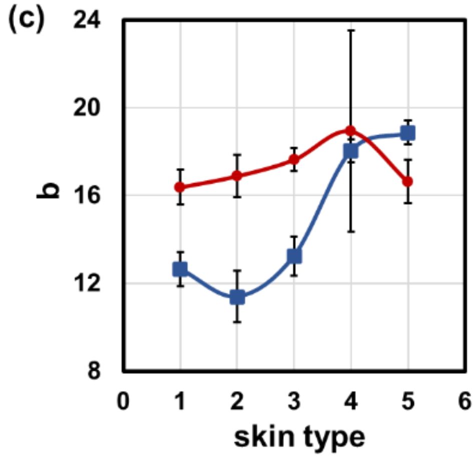


(d)

**Correlation Coefficients**

| | *L** | *a** | *b** |
|---|---|---|---|
| Volar, MVF | -0.89 | 0.51 | 0.94 |
| Dorsal, MVF | -0.78 | 0.78 | 0.68 |

**Figure S1. Summary of colorimetry measurements.** (a, b, c) The lightness (L*), *a**, and *b** parameters from colorimetry measurements for each skin phototype. The data points and error bar in each plot correspond to the average and standard deviation of each measurement, respectively. (*N*=6). The blue and red curves correspond to volar and dorsal forearms, respectively. (d) The correlation matrix of the colorimetric parameters and the melanin volume fractions (MVF).

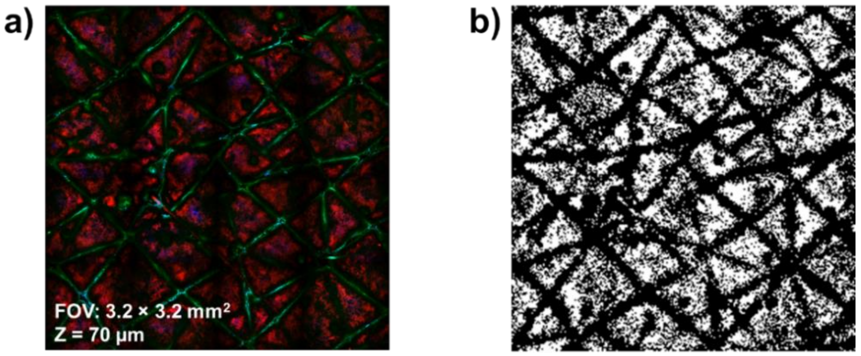


**Figure S2.** (a) An example of a raw 3-channel intensity images acquired in real-time with FLAME. (b) The equivalent melanin binary image obtained from the difference in the red and green channels, where thresholding was used to set pixels with difference in intensity > 0 to 1, and difference of ≤ 0 to 0.

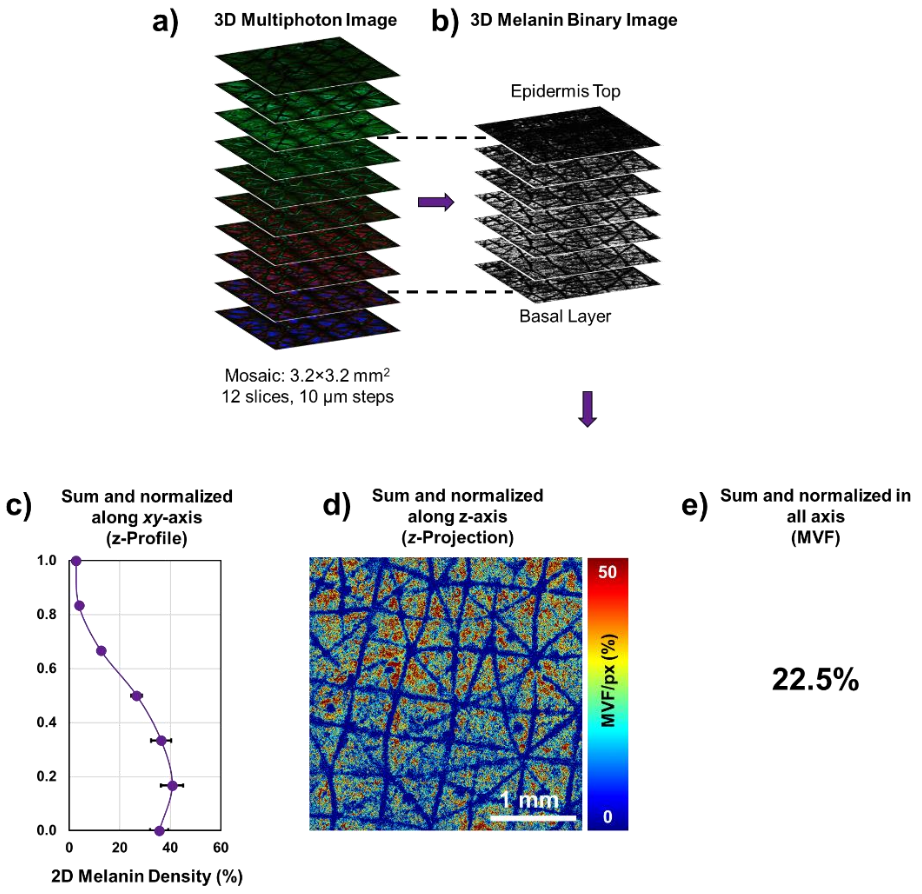


**Figure S3.** Schematic diagram for the data analysis. a) Raw 3-channel MPM volumetric image. b) The corresponding melanin binary volumetric image from the top of the epidermis to the basal layer. c) The resulting *z*-profile when the sum of (b) is taken along the *xy*-axis and normalized against the total number of pixels for each slice. d) The resulting *z*-projection when the sum of voxels in (b) is taken and normalized against the total number of pixels for each voxel. e) MVF is calculated by taking the global sum of (b) and divided by its total number of pixels.

**Table S1a.** Analysis of Variance (ANOVA) for the average global MVF of volar forearm

| Source | SS | df | MS | F | Prob > F |
|---|---|---|---|---|---|
| Columns | 2.215 | 4 | 0.554 | 147.140 | 1.044E-51 |
| Error | 0.583 | 155 | 0.004 | | |
| Total | 2.798 | 159 | | | |

**Table S1b**. *P*-values from Tukey-Kramer post-hoc analysis for ANOVA from Table S1a

| Type | II | III | IV | V |
|---|---|---|---|---|
| I | 4.389E-02 | 1.881E-06 | 0.000E+00 | 0.000E+00 |
| II | | 1.062E-01 | 1.239E-17 | 0.000E+00 |
| III | | | 4.158E-09 | 0.000E+00 |
| IV | | | | 0.000E+00 |

**Table S2a.** Analysis of Variance (ANOVA) for the average global MVF of dorsal forearm

| Source | SS | df | MS | F | Prob > F |
|---|---|---|---|---|---|
| Columns | 3.670 | 4 | 0.917 | 235.709 | 8.684E-65 |
| Error | 0.603 | 155 | 0.004 | | |
| Total | 4.273 | 159 | | | |

**Table S2b**. *P*- values from Tukey-Kramer post-hoc analysis for ANOVA from Table S2a

| Type | II | III | IV | V |
|---|---|---|---|---|
| I | 1.355E-08 | 8.104E-16 | 0.000E+00 | 0.000E+00 |
| II | | 1.861E-01 | 0.000E+00 | 0.000E+00 |
| III | | | 1.738E-14 | 0.000E+00 |
| IV | | | | 0.000E+00 |

**Table S3**. Summary of two-sample unpaired t-test for the comparison of average MVF from volar and dorsal forearm

| Type | Mean MVF ± S.D. (%) Volar | | | Mean MVF ± S.D. (%) Dorsal | | | *P*-value |
|---|---|---|---|---|---|---|---|
| I | 6.78 | ± | 6.32 | 9.67 | ± | 9.67 | 1.637E-01 |
| II | 11.04 | ± | 3.41 | 19.12 | ± | 3.41 | 1.752E-05 |
| III | 14.77 | ± | 4.03 | 22.52 | ± | 4.03 | 1.307E-11 |
| IV | 24.35 | ± | 5.79 | 34.84 | ± | 5.79 | 7.128E-12 |
| V | 39.83 | ± | 6.31 | 53.59 | ± | 6.31 | 2.175E-11 |

**Table S4a.** Analysis of Variance (ANOVA) for the measurements shown in Figure 5 in main text

| Source | SS | df | MS | F | Prob > F |
|---|---|---|---|---|---|
| Columns | 26.39 | 2 | 13.1928 | 0.8702 | 0.4317 |
| Error | 363.86 | 24 | 15.1608 | | |
| Total | 390.24 | 26 | | | |

**Table S4b***. P*- values from Tukey-Kramer post-hoc analysis for ANOVA from Table S1a

| Trial | T2 | T3 |
|---|---|---|
| T1 | 0.475 | 0.996 |
| T2 | | 0.524 |

**Table S5.** Sample size estimate as a function of FOV size for the detection of 10 to 25% change in the MVF values based on two-sample unpaired t-test (Power: 80% and significance level: 5%)

| FOV ($mm^2$) | Mean ± S.D. (%) | | | 10% | 15% | 20% | 25% |
|---|---|---|---|---|---|---|---|
| $0.25^2$ | 53.6 | ± | 11.6 | 74 | 33 | 19 | 12 |
| $0.29^2$ | 53.6 | ± | 10.2 | 57 | 26 | 15 | 10 |
| $0.46^2$ | 53.6 | ± | 7.7 | 33 | 15 | 9 | 6 |
| $0.81^2$ | 53.6 | ± | 6.3 | 22 | 10 | 6 | 4 |
| $1.62^2$ | 53.6 | ± | 5.0 | 14 | 7 | 4 | 3 |
